# Calcium Signaling in Oligodendrocyte Precursor Cells Mediated by Spontaneous and Evoked Responses: A Modeling Investigation

**DOI:** 10.1101/2025.08.14.670262

**Authors:** Martin Lardy, Leqi Wang, Claire Guerrier, Veronica T. Cheli, Pablo M. Paez, Anmar Khadra

## Abstract

Calcium (Ca^2+^) signaling has emerged as a central regulator of activity-dependent myelination in oligodendrocytes. These Ca^2+^ signals encompass both the stimulus-independent spontaneous Ca^2+^ local transients (SCaLTs) generated intrinsically in a voltage-independent manner or facilitated by the membrane voltage, as well as evoked responses triggered by ATP and glutamate release. To investigate the regulatory mechanisms underlying this combined spiking activity, we developed a stochastic spatiotemporal flux-balance model of Ca^2+^ transients in oligodendrocyte precursor cells (OPCs). The model incorporates all the relevant fluxes in these cells and integrates membrane voltage dynamics with a Ca^2+^-induced Ca^2+^-release (CICR) mechanism using parameters fitted to Ca^2+^ fluorescence recordings. The model reproduced the intrinsic and voltage-facilitated SCaLTs in OPCs in the absence of purinergic and glutamatergic receptors, and captured the three distinct patterns of evoked Ca^2+^ responses induced by ATP and glutamate identified using machine classifier. The model highlighted the role of ATP and glutamate concentrations in generating these clusters, and showed that the fast dynamics of CICR is key to producing these evoked responses. Further analysis of the model also revealed that voltage-gated L- and T-type Ca^2+^ channels slightly increase the frequency of SCaLTs, while stimulation with ATP and glutamate, using randomly distributed pulses mimicking in vivo conditions, leads to an increase in both the amplitudes of Ca^2+^ spikes (i.e., the combination of SCaLTs and evoked responses) and the prevalence of wide spikes, especially upon glutamate stimulation. Bifurcation analysis of the deterministic version of the model, in the absence of diffusion, demonstrated that ATP and glutamate stimulation can shift the system into an oscillatory regime, thereby increasing the deterministic component of SCaLT dynamics. This study thus offers a comprehensive representation of OPC Ca^2+^ transients linking recorded *in vitro* behaviors to *in vivo* dynamics.

**Author summary:** Oligodendrocytes are glial cells in the central nervous system that form myelin, the insulating sheath enabling rapid nerve signal transmission. Myelination is a dynamic process influenced by neuronal activity, with calcium (Ca^2+^) signaling emerging as a key regulator. These signals include spontaneous local Ca^2+^ transients (SCaLTs), generated intrinsically or facilitated by membrane voltage, as well as evoked responses triggered by neurotransmitters like ATP and glutamate. To understand how these signals arise and interact, we combined experimental recordings of Ca^2+^ activity in oligodendrocyte precursor cells (OPCs) with a data-driven biophysical model. The model incorporates stochastic Ca^2+^ fluxes, membrane voltage dynamics, and ca-induced Ca^2+^-release (CICR), allowing us to simulate diverse patterns of Ca^2+^ transients. Our simulations reproduced both intrinsic and voltage-facilitated SCaLTs and captured three distinct evoked response types induced by ATP and glutamate. We found that voltage-gated Ca^2+^ channels slightly enhance SCaLT frequency, while rapid CICR dynamics are critical for shaping the amplitude and timing of evoked signals. Furthermore, neurotransmitter stimulation can drive the system into an oscillatory regime, increasing the deterministic structure of Ca^2+^ transients. This work offers a mechanistic framework linking intracellular Ca^2+^ dynamics to the regulation of activity-dependent myelination in OPCs.

## 1 INTRODUCTION

Oligodendrocyte progenitor cells (OPCs) represent a distinctive type of stem cells within the central nervous system (CNS) of vertebrates, including humans [1]. Their significance stems from their crucial role in both the development and ongoing maintenance of the nervous system [2]. As OPCs mature, they undergo differentiation into oligodendrocytes, the cells that synthesize myelin, a lipid-rich substance, that ensheathe axons and support saltatory conduction by significantly enhancing the speed and efficiency of neural signal transmission [3, 4, 5].

The intricate interplay between axons and myelinating oligodendrocytes is fundamentally orchestrated by ion homeostasis mechanisms localized at the interface between the myelin sheath and the axons [6]. A manifestation of this intricate communication is the regulation of cytosolic calcium (Ca^2+^) concentration, denoted as [Ca^2+^]_*i*_, within oligodendrocytes [7, 8]. The dynamics of [Ca^2+^]_*i*_ within the cell hold the potential to exert influence over myelin formation, remodeling, and other yet-to-be-fully-understood functions [9]. Remarkably, it was shown that myelin sheath elongation is facilitated by a high frequency Ca^2+^ transients and obstructed by buffering [10].

Spontaneous Ca^2+^ local transients (SCaLTs) have been observed in isolated OPCs [11]. These events arise from predominantly stochastic (random) processes modulated by underlying deterministic patterns [12]. SCaLTs are attributed to both intrinsic cellular mechanisms as well as to fluctuations in membrane voltage. A growing body of evidence highlights the role of key Ca^2+^ fluxes in shaping SCaLTs driven by intrinsic dynamics. Prominent among these fluxes are (i) store-operated Ca^2+^ (SOC) channels that rely on the three core proteins: ORAI1, STIM1, and STIM2 to regulate Ca^2+^ influx across the cell membrane in response to changes in Ca^2+^ concentration within the endoplasmic reticulum (ER) [13, 11], denoted by [Ca^2+^]_*ER*_, (ii) the sodium/calcium exchanger (NCX) that exchanges one Ca^2+^ ion for two sodium (Na^+^) ions [14], (iii) Ca^2+^ release events emanating from the ER via inositol-triphosphate receptors (IP3Rs) and Ryanodine receptors (RyRs), both of which are involved in Ca^2+^-induced Ca^2+^-release (CICR) [15, 16], (iv) Ca^2+^ efflux mediated by sarco/endoplasmic reticulum Ca^2+^-ATPase (SERCA) pumps [17], and plasma membrane Ca^2+^-ATPase (PMCA) pumps [18].

The synergy of these fluxes forms the basis for SCaLTs in OPCs. Indeed, previous modeling studies of Ca^2+^ signaling in these cells have shown that fluxes through SOC, NCX, CICR (via IP3Rs and RyRs), and PMCA and SERCA pumps are sufficient for producing intrensic SCaLTs [12]. Interestingly, these cells also express voltage-gated Ca^2+^ channels (VGCCs), including L- and T-type Ca^2+^ channels on their membrane [19, 20, 21], which facilitate Ca^2+^ influx in response to membrane depolarization, contributing to cellular signaling and function including migration and myelination [22, 21, 19]. Despite their importance, it remains unclear how voltage membrane fluctuations and Ca^2+^ entry through VGCCs affect SCaLTs and their underlying dynamics.

A pivotal aspect of OPC function is their ability to receive glutamatergic signals via *α*-amino-3-hydroxy-5-methyl-4-isoxazolepropionic acid receptors (AMPAR) and N-methyl-D-aspartate receptors (NMDAR) [23], [24]. These receptors mediate cation influx, including Ca^2+^, enabling glutamate signaling to elevate [Ca^2+^]_*i*_ either directly or indirectly [9] and to subsequently modulate subcellular processes such as proliferation and differentia-tion [25]. Additionally, the presence of purinergic P2X7 receptors (P2XRs) in OPCs has been implicated in diverse signaling pathways [9, 26, 27]. In mouse OPCs, P2X7R activation by ATP induces distinct signaling responses reminiscent of these previously analyzed [28, 29, 30, 31, 32], further underscoring its functional importance [9, 33]. However, the interplay between purinergic and glutamatergic receptors, along with membrane depolarization, in shaping evoked Ca^2+^ responses, particularly in the context of SCaLTs, remains an open area of investigation.

In this study, we extended a previously developed stochastic spatiotemporal flux-balance model,implemented in one spatial dimension and validated against fluorescent Ca^2+^ imaging data from rat OPCs [12], to investigate the dynamics of both spontaneous and evoked Ca^2+^ transients in OPCs. The results demonstrated the model’s ability to replicate key features of evoked Ca^2+^ responses observed *in vitro* while predicting how these responses manifest *in vivo*. Specifically, it delineated the distinct effects of membrane depolarizations, glutamate signaling and ATP stimulation, with the two latter ones modeled as random processes. Additionally, the model provided novel insights into the interplay between SCaLTs and evoked responses in OPCs, identifying the parameter regimes that drive these oscillatory behaviors.

## 2 METHODS

### 2.1 Experimental Methods

#### 2.1.1 Primary cultures of OPCs

Primary cultures of cortical OPCs were prepared as described [34, 35, 36] which results in >98% OPCs and <1% GFAP stained astrocytes or Iba1 stained microglia. Cerebral hemispheres from 1-day old mice were mechanically dissociated, then plated in poly-D-lysine-coated flasks in DMEM/F12 (1:1 v/v) (Invitrogen) supplemented with 10% fetal bovine serum (FBS) (Life Technologies). After 4 h, the medium was changed and cells grown in DMEM/F12 supplemented with insulin (5 *µ*g/ml), apotransferrin (50 *µ*g/ml), Na^+^ selenite (30 nM), d-Biotin (10 mM) and 10% FBS (Life Technologies). Every 3 days 2/3 of the media was changed. OPCs were purified from mixed glia after 14 days by a differential shaking and adhesion procedure. The detached cells were plated into Petri dishes for 30 min at 37°C to allow microglia and astrocytes to adhere, then non-attached cells collected and plated on poly-D-lysine-coated coverslips in DMEM/F12 supplemented with insulin (5 *µ*g/ml), apotransferrin (50 *µ*g/ml), Na^+^ selenite (30 nM), 0.1% BSA, progesterone (0.06 ng/ml), putrescine (16 *µ*g/ml) (Sigma), and 2% FBS. OPCs were kept in mitogens, platelet derived growth factor (PDGF) and basic fibroblast growth factor (bFGF, 20 ng/ml) (Peprotech), for 3 days.

#### 2.1.2 Ca^2+^ imaging

Primary cultures of cortical OPCs were prepared as described above. Before imaging, OPCs were washed in serum and phenol-red-free DMEM and incubated for 25 min with 4 *µ*M fura-2 (AM) (Life Technologies) plus 0.08% Pluronic F127 (Life Technologies) at 37°C and 5% CO_2_. Cells were then washed four times in DMEM and stored in DMEM for 10 min before being imaged. Ca^2+^ influx and resting Ca^2+^ levels were measured in serum and phenol-red-free HBSS containing 1.3 mM Ca^2+^ and 1 mM Mg^2+^. Fura-2 was excited by alternating 340 and 380 nm. Fluorescence signals were acquired every 2 s by means of a high-speed wavelength-switching device (Sutter Instruments, Lambda DG4). A spinning disc confocal inverted microscope (Olympus, IX83-DSU) equipped with a CCD camera (Hamamatsu, ORCA-R2) measured fluorescence. Ca^2+^ influx and resting Ca^2+^ levels were measured on individual cell bodies using the image analysis software Meta Fluor (Molecular Devices). To minimize bleaching, excitation light intensity and sampling frequency was kept as low as possible.

### 2.2 Mathematical Methods

#### 2.2.1 Mathematical model

A stochastic spatiotemporal model describing the dynamics of intrinsic (or voltage-independent) SCaLTs in OPCs in one spatial dimension was previously developed [12]. We extended this model in the current study to account for evoked Ca^2+^ responses, which also entails considering the concentrations of various cations inside and outside the OPCs, including Ca^2+^, Na^+^ and potassium (K^+^) (Fig. 1A-C). Two sets of fluxes were incorporated into the model (Fig. 1A): (i) Those occurring at the cell membrane, through voltage gated Ca^2+^ channels (VGCC) including L-type and T-type Ca^2+^ channels (*J*_*L*−*type*_ and *J*_*T*−*type*_, respectively), store operated channels (*J*_*SOC*_), glutamatergic AMPARs and NMDARs (*J*_*AMPA*_ and *J*_*NMDA*_, respectively), purinergic P2X7Rs (*J*_*P*2*X*7_), Na^+^/Ca^2+^ exchangers (*J*_*NCX*_), Na^+^/K^+^ exchangers (*J*_*NaK*_), K^+^ leak (*J*_*KLeak*_) and PMCA pumps (*J*_*PMCA*_). (ii) Those occurring at the ER membrane, through IP3Rs (*J*_*IP*3_), RyRs (*J*_*Ry*_), Ca^2+^ leak (*J*_*Leak*_) and SERCA pumps (*J*_*SERCA*_).

**Figure 1.**
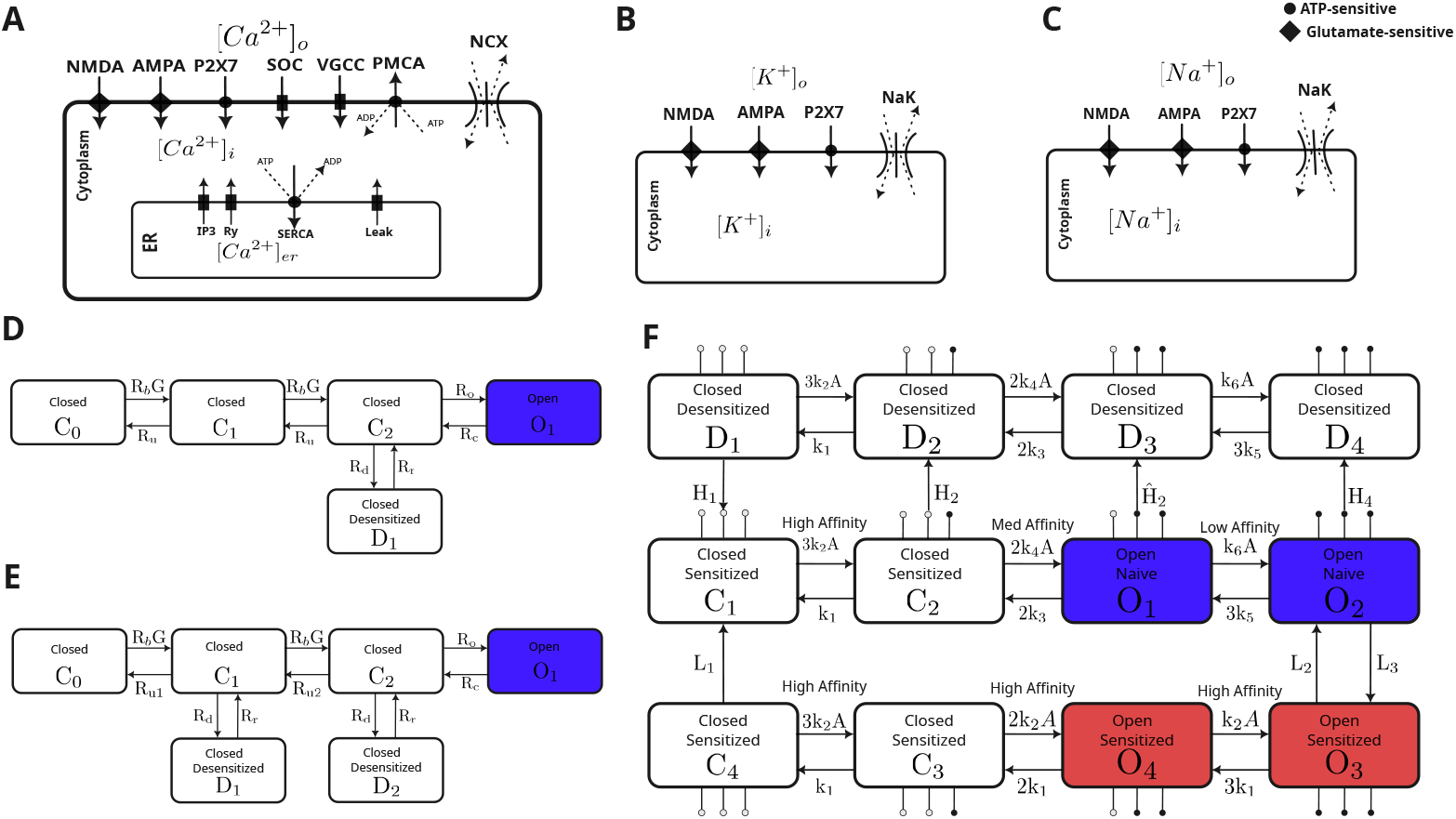
Schematic of the SMM and receptor-specific kinetic Markov models integrated within the SSM. (A) Diagram of the cell along with the Ca^2+^ fluxes across the plasma and ER membranes. (B) Diagram of the cell along with the K^+^ fluxes across the plasma membrane. (C) Diagram of the cell along with the Na^+^ fluxes across the plasma membrane. (D) Kinetics of the NMDAR. (E) Kinetics of the AMPAR. (F) Kinetics of the P2X7R. The kinetics of the three receptors in D, E and F are described in terms of the states *C*_*i*_ representing the non-conducting closed states (*i* = 0, 1, 2 for NMDAR and AMPAR, and *i* = 1, 2, 3, 4 for P2X7R), *D*_*i*_ representing the non-conducting desensitized states (*i* = 1 for NMDAR, *i* = 1, 2 for AMPAR and *i* = 1, 2, 3, 4 for P2X7R) and *O*_*i*_ representing the conducting open (blue) and sensitized (red) states (*i* = 1 for NMDAR and AMPAR, and *i* = 1, 2, 3, 4 for P2X7R), with transitions rates: *R*_*b*_, *R*_*u*_, *R*_0_, *R*_*c*_, *R*_*d*_, *R*_*r*_ for NMDAR, *R*_*b*_, *R*_*u*_, *R*_0_, *R*_*c*_, *R*_*d*_, *R*_*r*_ for AMPAR and *k*_*j*_, *H*_*m*_, *L*_*n*_ (*j* = 1, … 6, *m* = 1, 2 and *n* = 1, 2, 3) for P2X7R, where *G* = [Glut] and *A* = [ATP] are the concentrations of glutamate and ATP, respectively.

The resulting extended model was formulated using a set of stochastic ordinary and partial differential equations that capture the dynamics of cytosolic Ca^2+^ ([Ca^2+^]_*i*_), ER Ca^2+^ ([Ca^2+^]_*ER*_), as well as cytosolic Na^+^ ([Na^+^]_*i*_) and K^+^ ([K^+^]_*i*_) concentrations. Diffusion was explicitly incorporated into the equations governing the cytosolic concentrations of these cations as described below

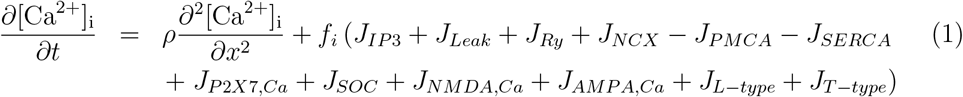

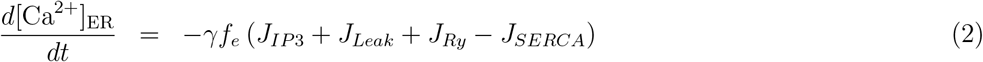

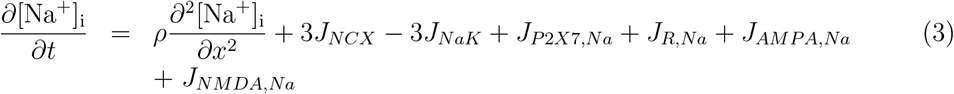

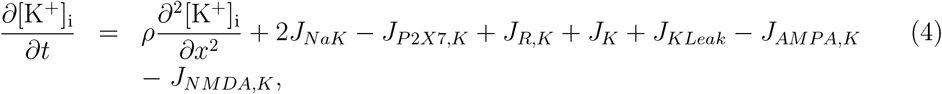

where *J*_*P*2*X*7,*X*_, *J*_*NMDA,X*_, *J*_*AMPA,X*_ are the contributions of P2X7R, NMDAR and AMPAR, respectively, to the influx of Ca^2+^ (*X* = *Ca*) and Na^+^ (*X* = *Na*), and efflux of K^+^ (*X* = *K*), whereas *ρ* is the diffusion coefficient of all ions.

The Li-Rinzel model for IP3R kinetics [37] was used to describe *J*_*IP*3_ that depends on both [Ca^2+^]_*i*_ and cytosolic IP3 concentration [IP3]_*i*_. Specifically,

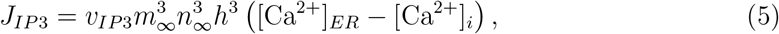

Where

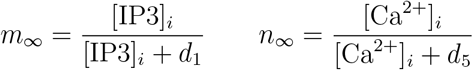

and the gating variable *h* satisfies

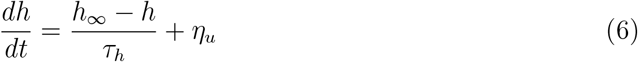

With

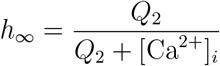

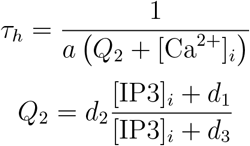

and [IP3]_*i*_ satisfies the equation

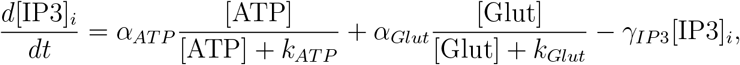

where [ATP] and [Glut] are the concentrations of ATP and glutamate applied. The term *η*_*u*_ in Eq. (6) is an Ornstein-Uhlenbeck noise process that satisfies the equation

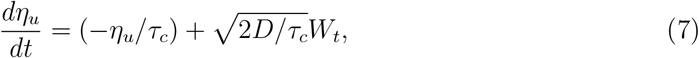

where *τ*_*c*_ is the characteristic correlation time of the noise, *D* is the noise intensity and *W*_*t*_ is a Gaussian white noise process with *µ* = 0 and *σ* = 1 [12]. This noise was used to produce the slow baseline oscillations in Ca^2+^ due to the stochastic slow clustering of IP3Rs [12].

The flux through RyRs (*J*_*Ry*_) was described according to the Levine-Keizer model [38], given by

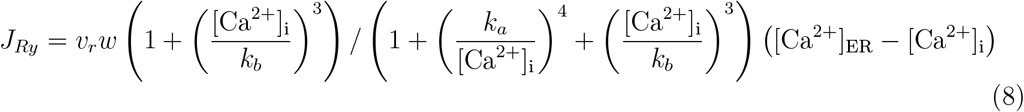

where *w* is a slow gating variable that satisfies

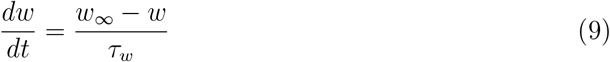

With

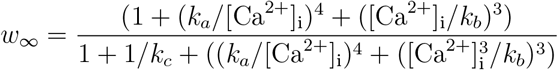

And

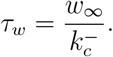

A Hill function was used to describe the fluxes through the PMCA and SERCA pumps [39], according to the equations

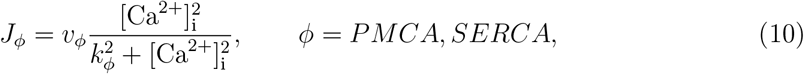

whereas [Na^+^]_*i*_ and and [K^+^]_*i*_ were assumed to return to rest according to the equations

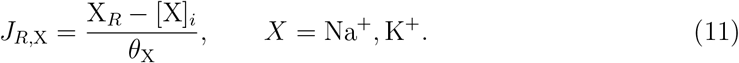

Here 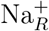 and 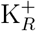 represent the resting concentrations of Na^+^and K^+^, respectively, while *θ*_*Na*_ and *θ*_*K*_ are the time constants of the concentrations of these two ions to return to rest.

To define the remaining fluxes in Eqs. (1)-(4), we first need to introduce the equation for membrane voltage using the Hodgkin-Huxley formalism [40], given by

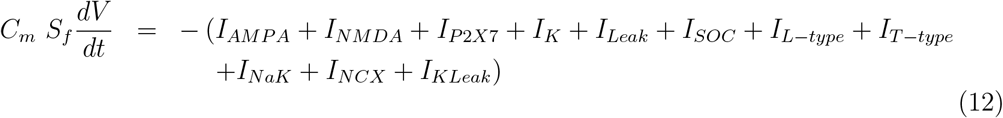

where *I*_*Y*_ (*Y* = *AMPA, NMDA, P*2*X*7, *K, Leak, SOC, L*−*type, T*−*type, NaK, NCX, KLeak*) are the various ionic currents produced by ion flow through channels and exchangers expressed on the membrane of OPCs, *S*_*f*_ is the membrane surface and *C*_*m*_ is the specific membrane capacitance. The fluxes associated with these currents (including those for the two VGCCs: *I*_*L*−*type*_ and *I*_*T*−*type*_) were then computed using the equation [39]

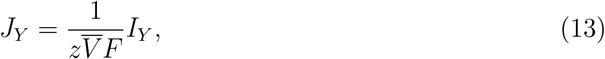

where *z* is the valence, 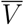 is the cell volume, and *F* is Faraday’s constant.

We used Destexhe et al. NMDAR and AMPAR models [41] (Fig. 1D and E, respectively) to describe *I*_*NMDA*_ and *I*_*AMPA*_, given by

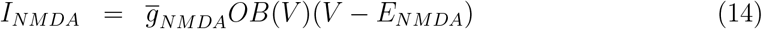

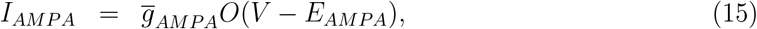

where *O* is the open state of each receptor, *E*_*Y*_ and 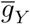 are, respectively, the Nernst potential and maximum conductance of NMDARs (*Y* = *NMDA*) and AMPARs (*Y* = *AMPA*), and *B*(*V*) is the voltage-dependent magnesium (Mg^2+^) block, given by

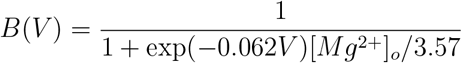

where [Mg^2+^]_*o*_ is the extracellular Mg^2+^ concentration.

Using the Khadra et al. P2X7R model [29] to describe *I*_*P*2*X*7_ (Fig. 1F), we have

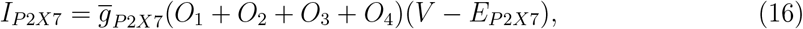

where *O*_1_, *O*_2_ are the open states and *O*_3_, *O*_4_ are the sensitized states of P2X7R, 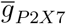 is the receptor maximum conductance and *E*_*P*2*X*7_ is its Nernst potential (set to zero because P2X7Rs are nonspecific).

For the inward-rectifying K^+^ current (*I*_*K*_), we adopted the formalism of [42, 43] to describe it, i.e.,

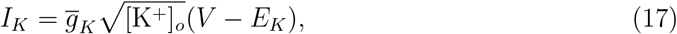

where [K^+^]_*o*_ is the extracellular K^+^ concentration, 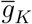 is the maximum conductance of *I*_*K*_, and *E*_*K*_ is the K^+^ Nernst potential that depends on both the gas constant (*R*) and absolute temperature 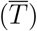 as follows

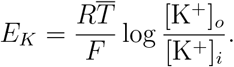

Similarly, we adopted the formalisms from [44] and reparameterized them using the data from [20] to describe the two VGCCs: L-type and T-type Ca^2+^ currents (*I*_*L*-*type*_ and *I*_*T*-*type*_, respectively) included in the SSM Fig. 2). In this case,

**Figure 2.**
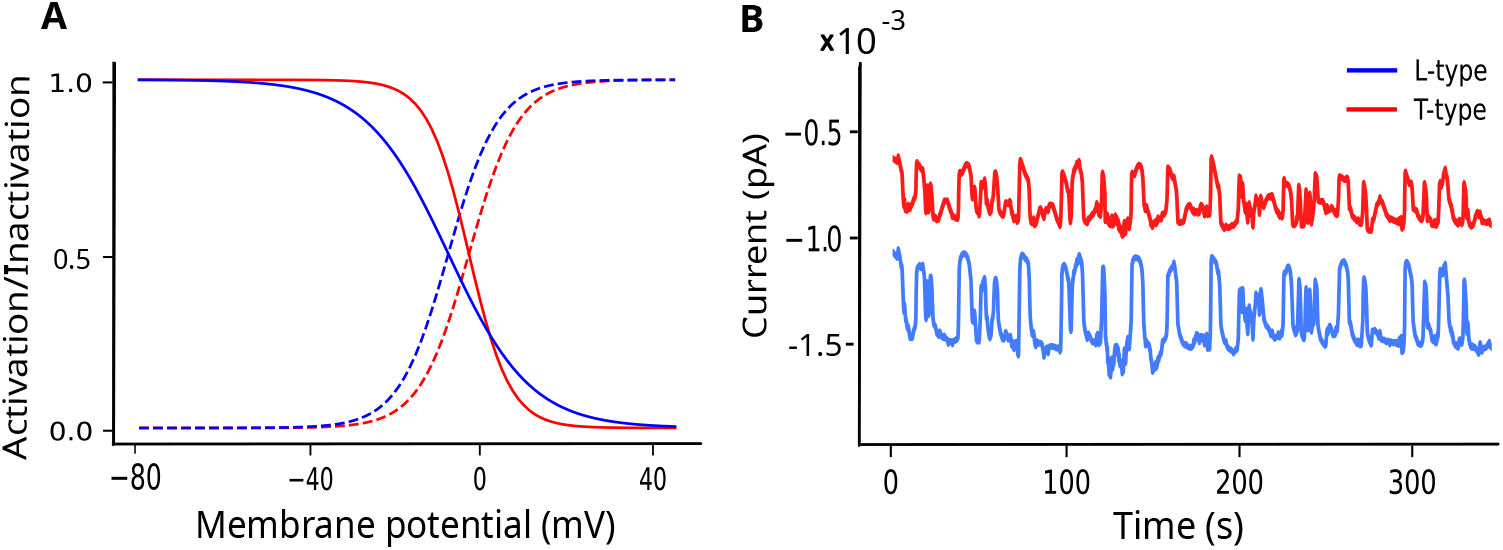
Kinetics of the L-type and T-type Ca^2+^ channels. (A) The steady state activation and inactivation curves of the L-type and T-type Ca^2+^ channels color-coded according to the legend. (B) The stochastic current fluctuations produced by the L-type and T-type Ca^2+^ channels color-coded according to the legend.

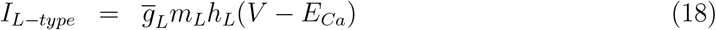

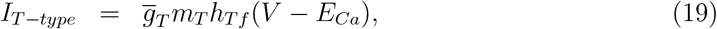

where *E*_*ca*_ is Ca^2+^ Nernst potential, given by

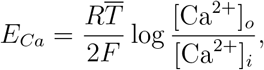

and 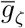 is the maximum conductance of *I*_*L*_ (*ζ* = *L*) and *I*_*T*_ (*ζ* = *T*). The two gating variables *m*_*L*_ and *h*_*L*_ are the activation and inactivation variables of *I*_*L*_, respectively; their dynamics are governed by the equations

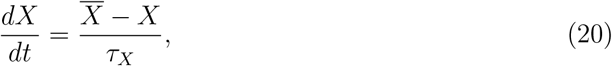

where 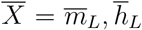 are the steady state activation and inactivation, respectively, and 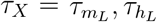 are the time constants, given by

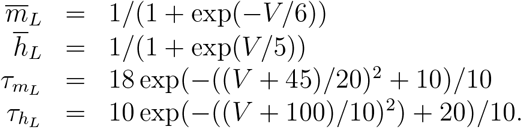

The gating variables of the current *I*_*T*_, on the other hand, are the activation variable *m*_*T*_ and the inactivation variable *h*_*Tf*_; their dynamics are governed by Eq. (20), where 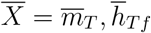 are the steady state activation and inactivation, respectively, and 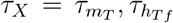 are the time constants, given by

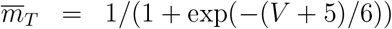

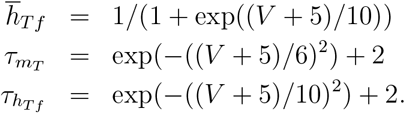

The SOC current (*I*_*SOC*_) expressed in [45] was adopted in this study; its expression is given by

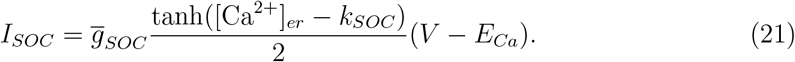

Two leak currents were included in the voltage equation (12): a K^+^ leak current (*I*_*KLeak*_) and a standard leak current (*I*_*Leak*_). These two currents are given by

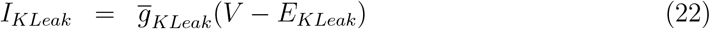

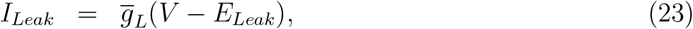

where *E*_*KLeak*_, *E*_*Leak*_ are Nernst potential of these currents, whereas *g*_*KLeak*_ and *g*_*Leak*_ are their conductances. Na^+^ leak was not included in the model because it was assumed to be small.

Finally, two exchangers were included in the model, the Na^+^/Ca^2+^ exchanger (*I*_*NCX*_) and the K^+^/Na^+^ exchanger (*I*_*NaK*_). Using the formalisms for these two currents introduced in [46], we have

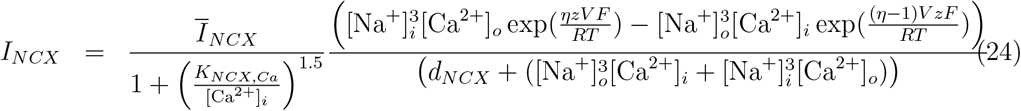

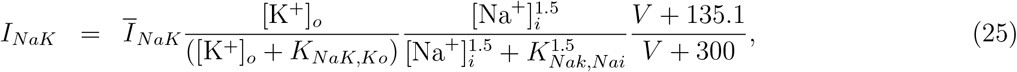

where *η* is the position of the energy barrier, *Ī*_*NCX*_ and *Ī*_*NaK*_ are the maximum currents generated by Na^+^/Ca^2+^ and K^+^/Na^+^ exchangers, respectively, and *K*_*NCX,Ca*_, *d*_*NCX*_, *K*_*NaK,Ko*_ and *K*_*NaK,Nai*_ are parameters representing half-maximum activation.

Parameter values of this stochastic spatiotemporal model (SSM), given by Eqs. (1)-(25), and their definitions are provided in Table 2.2.2. By ignoring noise, i.e., removing Eq. (7), and by setting diffusion to zero (i.e., *ρ* = 0), we obtain the deterministic temporal model (DTM). In the absence of membrane voltage dynamics and the kinetics of P2X7Rs, AMPARs and NMDARs, the SSM was able to reproduce the original features of the model associated with intrinsic ScaLTs previously reported [12] (Fig. S1).

#### 2.2.2 Random ATP and glutamate stimulations

The SSM model Eqs. (1)-(25) was subjected to random ATP and glutamate stimulations. These random stimulations were modeled as spike trains with frequencies following a binomial distribution ℬ(*n, p*), where *n* = 1 and *p* = 0.001, 0.0025, and 0.005 corresponding to low, medium, and high spike frequencies, respectively. Random sampling from this distribution was performed every 1 ms of simulation time, resulting in average interspike intervals of 10, 4, and 2 s, respectively. The amplitudes of the ATP and glutamate spike trains followed a norma distribution 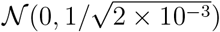, leading to an overall spike distribution of 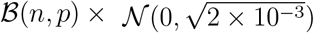. To analyze the simulated Ca^2+^ signals under each stimulation condition, we conducted 50 simulations, each lasting 300 s, and recorded Ca^2+^ transients exceeding 0.35 *µ*M. The threshold was established based on the mean Ca^2+^ concentration in wild-type simulations, corresponding to the default parameter values in Table 2.2.2. The amplitudes of collected Ca^2+^ transients were then binned to construct a distribution. Similarly, the width of these transients were quantified by computing their full width at half maximum (FWHM) and the results were binned to construct a distribution.

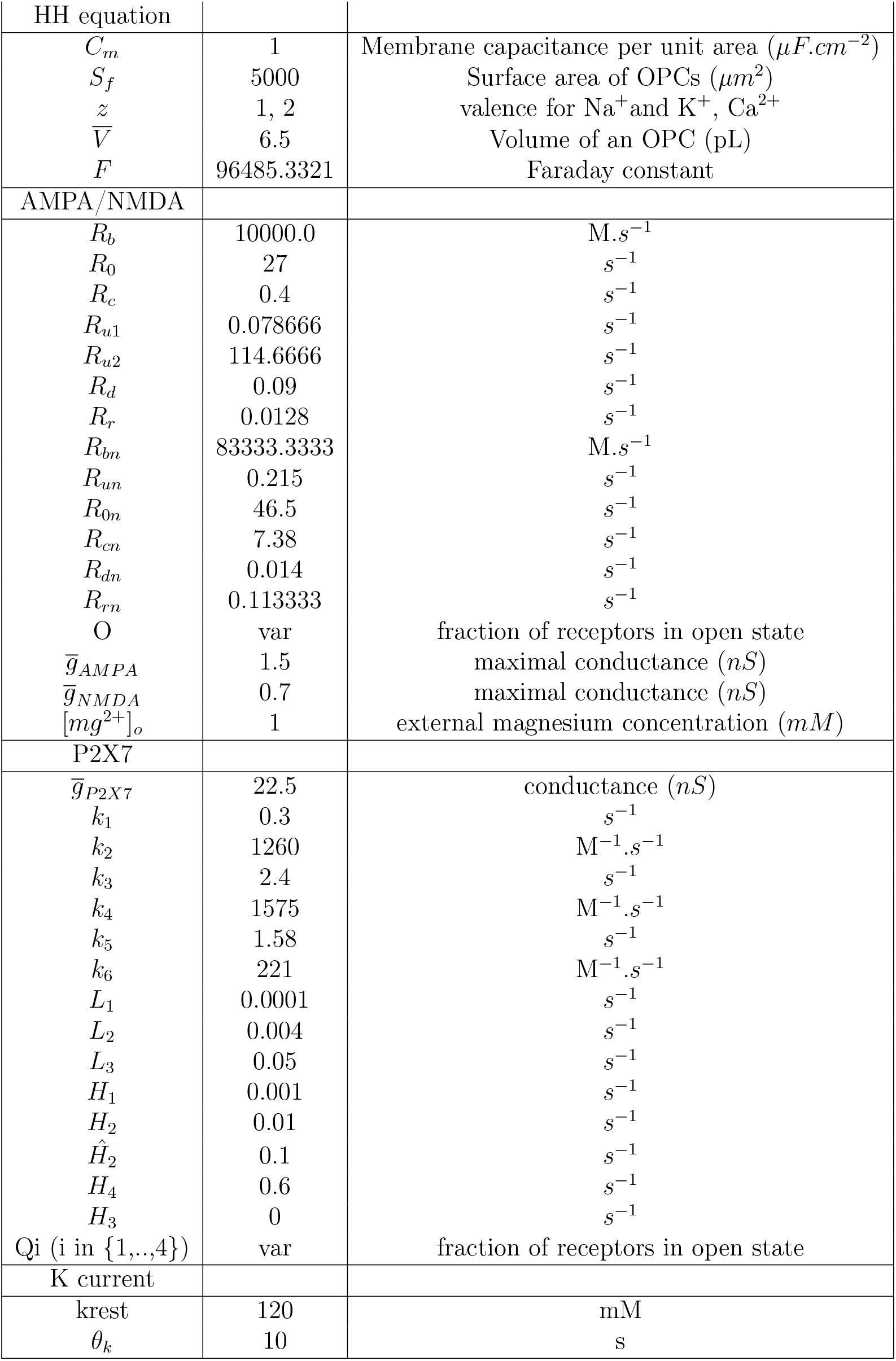
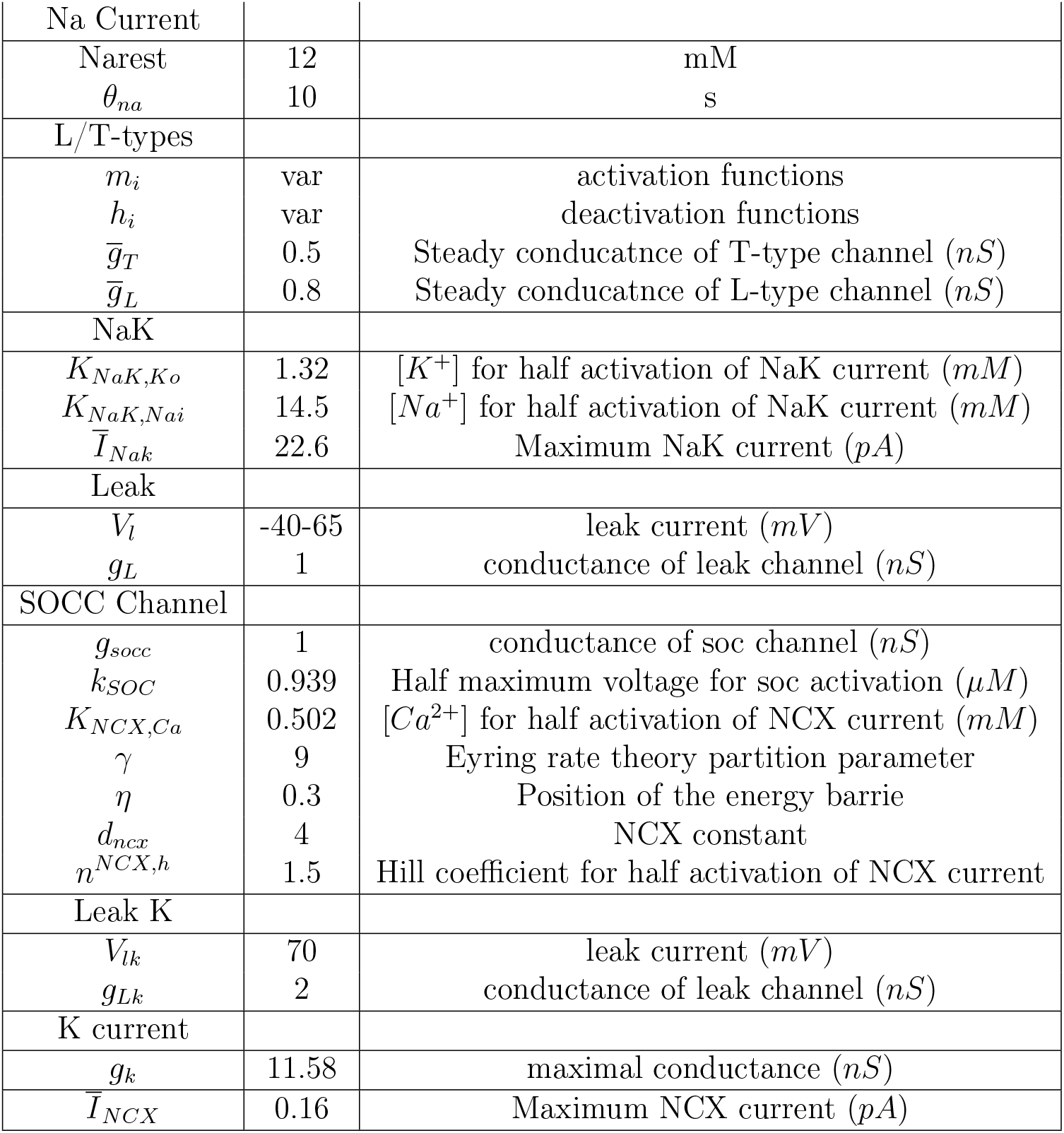

#### 2.2.3 Numerical methods and software

A forward time-centered space scheme with Neumann conditions at the boundary was used to compute numerical solutions to the SSM. A custom-made python script was used to compute these solutions. The DTM was numerically solved in XPPAUT (a freeware available online at http://www.math.pitt.edu/~bard/xpp/xpp.html) to compute bifurcation diagrams. Ca^2+^ transients were classified as Ca^2+^ spikes if they exceeded a threshold, defined as one standard deviation above the mean of the simulated signal generated by the SSM under default parameter values (also referred to as the WT model). To compare spikes across different conditions, such as the WT model versus perturbed versions where L- and T-type Ca^2+^ channels are blocked (a version that is referred to as the KO model), the average threshold from WT simulations was used as a reference for both conditions. To perform clustering analysis, each time series was fitted with a 4th-degree polynomial, and the resulting polynomial coefficients were used as input for a k-means algorithm. The optimal cluster number was determined to be three using the elbow score. Computer sim-ulations of the SSM and the DTM along with the clustering analysis were performed using a laptop with a IntelCore i7-8550U CPU at 1.80GH with 16,0Go of RAM. Version of the SSM solver code and the deterministic code for XPPAUT are available for download at https://github.com/martinLARD/SSM-model.git, respectively, in Python and Julia.

### 2.3 Quantification of Ca^2+^ Transients

SCaLTs (including those facilitated by the voltage) were isolated by eliminating Ca^2+^ fluxes associated with P2X7Rs, AMPARs and NMDARs from the SSM, as these fluxes are responsible for evoked responses. Following the approach in [12], Ca^2+^ transients, including SCaLTs and those influenced by fluxes involved in evoked responses, were identified by detecting peaks exceeding 20% above baseline or 1 standard deviation above the mean. To enable direct comparison between simulated and experimental data, the time series of both datasets were z-scored.

## 3 RESULTS

### Bifurcation Analysis of Evoked Responses

Incorporating the effects of ATP and glutamate concentrations ([ATP] and [Glut], respectively) into the model allows us to investigate how these exogenous stimuli, known to trigger evoked Ca^2+^ responses in OPCs, modulate the deterministic dynamics of the DTM. To this end, we performed bifurcation analyses of the membrane voltage (*V*) and cytosolic calcium concentration ([Ca^2+^]_i_) with respect to both [ATP] and [Glut], using the continuation method in XPPAUT.

In the case of [ATP], we found that both *V* (Fig. 3A) and [Ca^2+^]_i_ (Fig. 3C) exhibit three distinct regimes of behavior at low, intermediate and high [ATP]. At low and high [ATP], a stable branch of equilibria was observed, representing the quiescent state of the cell; these stable branches connect with each other through an unstable branch of equilibria for intermediate [ATP] at two Hopf bifurcations (HB1 to the left and HB2 to the right). Envelopes of stable limit cycles emerge from these two Hopf bifurcations and soon after become envelopes of unstable limit cycles when they undergo torus bifurcations (TR1 to the left and TR2 to the right), signifying the onset of chaotic dynamics. This latter regime corresponds to where evoked periodic responses in cytosolic Ca^2+^ induced by ATP can be observed. It is important to note that the quiescent regime to the left of HB1 is type III excitable allowing the model to generate SCaLTs reminiscent to those previously analyzed in [12].

**Figure 3.**
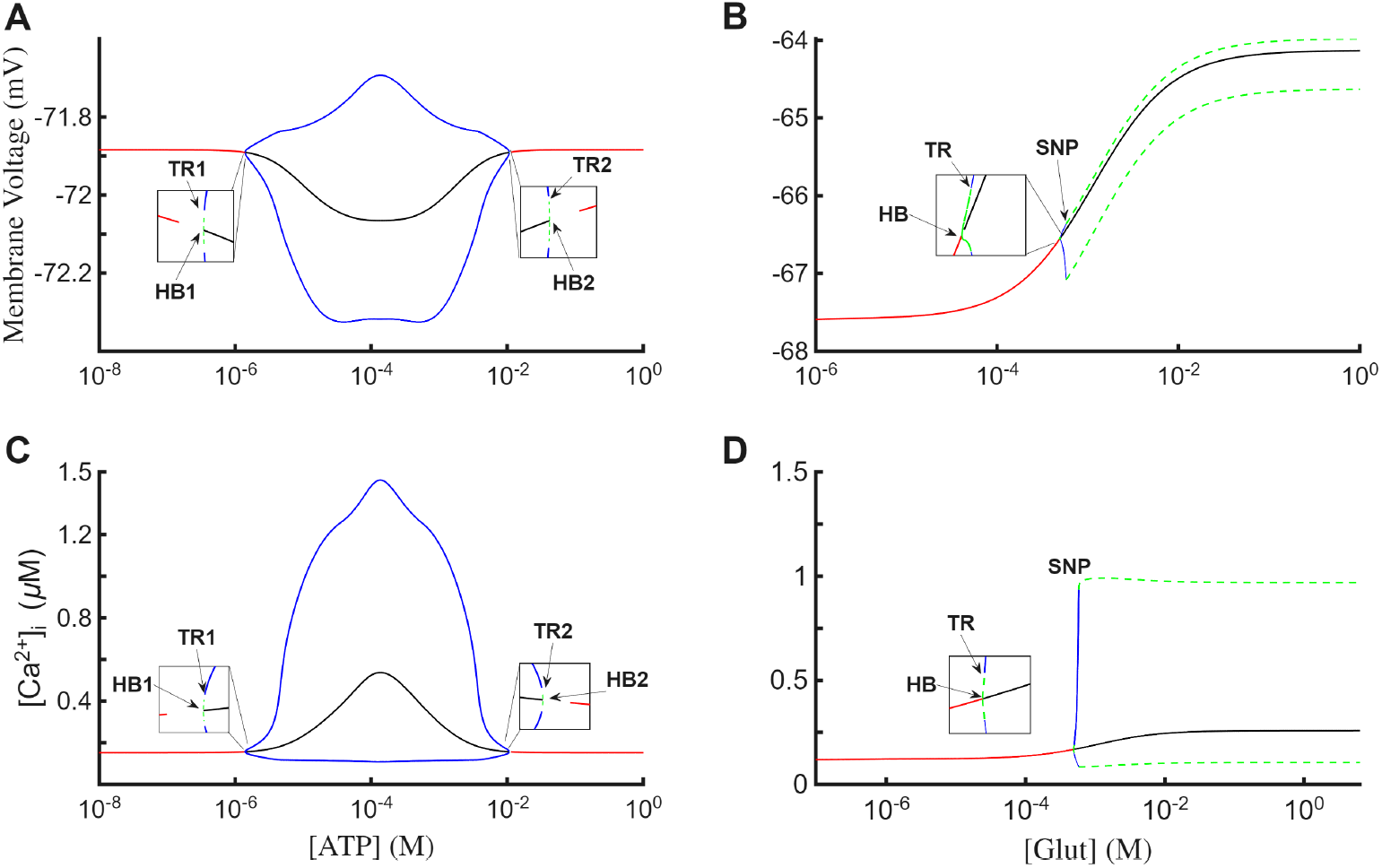
Steady state analysis of the DTM. Bifurcation diagrams of the membrane voltage (*V*) with respect to A) ATP concentration: [ATP], and (B) glutamate concentration: [Glut]. Bifurcation diagrams of cytosolic Ca^2+^ concentration ([Ca^2+^]_i_) with respect to (C) [ATP] and (D) [Glut]. Red (black) solid lines: Branches of stable (unstable) equilibria. Green (blue) solid lines: Envelopes of stable (unstable) limit cycles; the envelopes in A and C are delimited by two Hopf Bifurcations (HB1 to the left and HB2 to the right) and undergo two torus bifurcations (TR1 to the left and TR2 to the right), while the envelopes in B and D emerge from a Hopf bifucation (HB) and undergo both a torus bifurcation (TR) and a saddle-node of periodics bifurcation linking the stable and unstable envelopes.

By repeating the same analysis with [Glut], we also obtained three regimes of behavior for both *V* (Fig. 3B) and [Ca^2+^]_i_ (Fig. 3D): a quiescent regime at low [GLUT] defined by a stable branch of equilibria possessing type III excitability (allowing the model to produce SCaLTs), a mixed-mode oscillation regime at intermediate [GLUT] defined by envelopes of limit cycles that emerge from a Hopf bifurcation (HB) and immediately undergo torus bifurcation (TR) before terminating at a saddle-node of periodics (SNP) bifurcation, and a tonically spiking oscillatory regime at high [GLUT] formed by envelopes of stable limit cycles emerging from the SNP. The stable branch of equilibria at low [GLUT] regime losses stability at the HB and persists at intermediate and high [GLUT], while the periodic orbits in the mixed-mode oscillation regime combine both small and large amplitude oscillations within each cycle in a manner reminiscent to those previously observed [12, 47].

Together, these results demonstrate that both [ATP] and [Glut] can drive the system into regimes characterized by periodic or chaotic Ca^2+^ spiking, thereby accounting for evoked cytosolic Ca^2+^ responses in OPCs.

### Effects of L- and T-Type Ca^2+^ Channels on SCaLTs

It has been previously shown, using computational modeling that Ca^2+^-induced Ca^2+^-release (CICR) through IP3R and RyR, along with SOC and NCX fluxes on the plasma membrane and fluxes through SERCA and PMCA pumps, are sufficient for intrinsically generating SCaLTs in OPCs [12]. The flux balance model accounting for these fluxes successfully reproduced key features of Ca^2+^ signals, including SCaLT characteristics (e.g. slow baseline oscillations, doublets, random spiking, etc). This model, however, did not include the effects of VGCC expressed on OPC plasma membrane, a component that may have significant effects on SCaLTs.

To investigate this, two types of Ca^2+^ channels were included in the SSM, including the high-voltage activated L-type and low-voltage activated T-type Ca^2+^ channels. Stochastic simulation of the model showed that it continues to display random spiking events that surpass a threshold, defined as one standard deviation above the mean, reminiscent of SCaLTs (Fig. 4A); these spiking events integrate SCaLTs generated intrinsically with those facilitated by the voltage. To assess the contribution of VGCCs to SCaLTs, we compared the steadystate distribution of spike counts between WT (i.e., with L- and T-type Ca^2+^ channels included in the SSM) and KO (i.e., SSM lacking L- and T-type Ca^2+^ channels) conditions (Fig. 4B). Quantification was based on 50 steady-state simulations (60 s each), with the average WT threshold used to identify Ca^2+^ spikes in both conditions. Our results showed that VGCCs increase the number of spikes by expanding the tail of the distribution in WT compared to KO conditions. Further examination of the amplitude of all Ca^2+^ transients in these simulations, including sub- and supra-threshold ones, revealed that VGCCs increase not only the amplitude of spikes by generating a longer tail in WT condition (as highlighted by the red arrow in Fig. 4A), but also the amplitude of sub-threshold transients. These findings indicate that VGCCs further facilitate SCaLTs and contribute to the generation of more pronounced Ca^2+^ signals.

**Figure 4.**
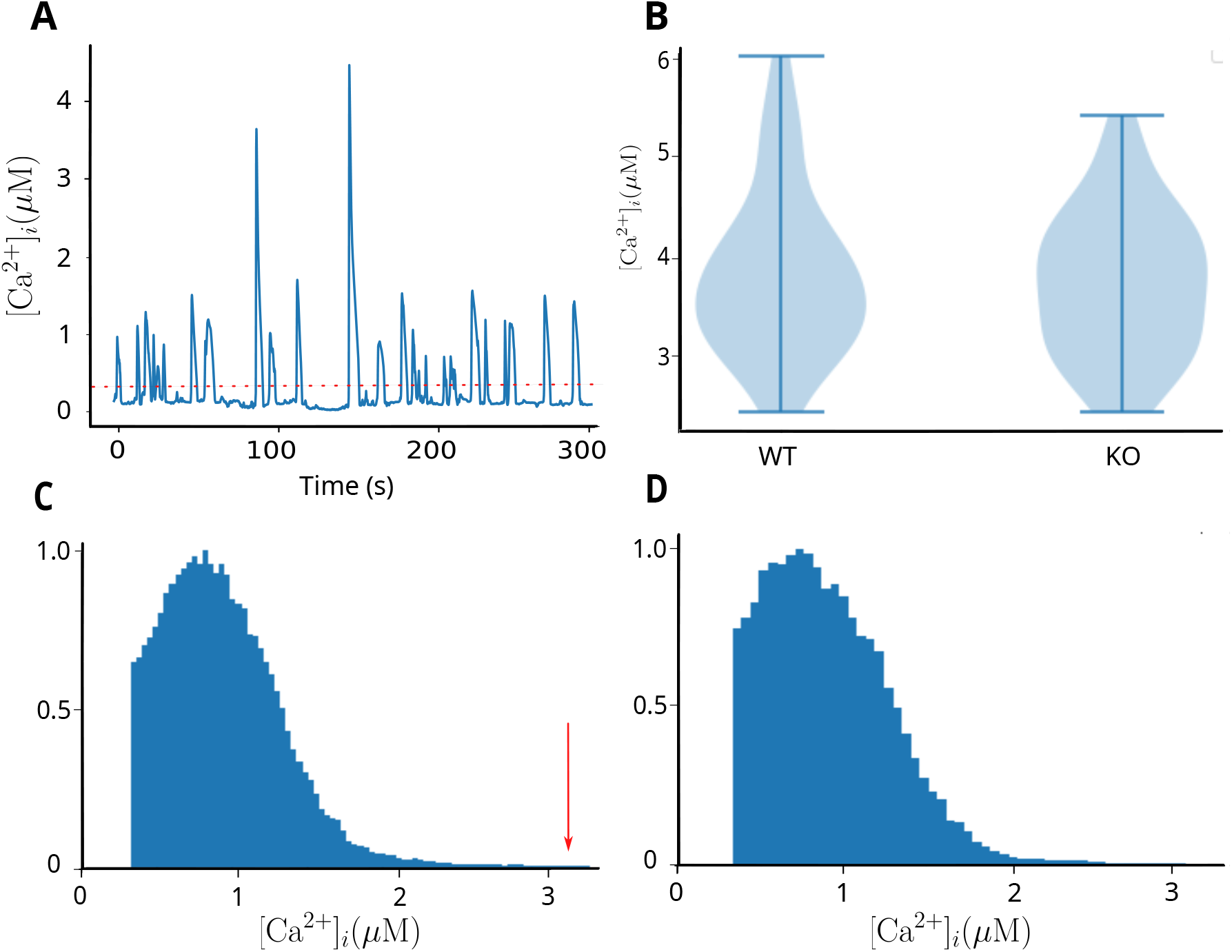
Contribution of L- and T-type Ca^2+^ channels to SCaLTs in OPCs as predicted by the SSM. (A) A simulated stochastic Ca^2+^ signal generated by the SSM displaying sub- and supra-threshold Ca^2+^ transients separated by a red line marking the mean. The suprathreshold VGCC-mediated Ca^2+^ transients combine the intrinsic and voltage-facilitated SCaLTs. (B) Violin plots of the number of SCaLTs generated by the SSM in the presence (labeled WT) and absence (labeled KO) of L- and T-type Ca^2+^ channels. A total of 50 steady state simulations of 60 s each were included per condition and the number of SCaLTs above the average threshold of WT condition were counted for both conditions. Note that SCaLTs in KO condition are essentially the voltage-independent (or intrinsic) SCaLTs. (C) Average amplitude distribution of sub- and suprathreshold Ca^2+^ transients for WT SSM obtained from the simulations in B. The red arrow indicates the extended tail in the WT condition. (D) Average amplitude distribution of sub- and supra-threshold Ca^2+^ transients for KO SSM obtained from the simulations in B.

### OPC Ca^2+^ Responses Induced by ATP

It has been previously shown that OPCs express ATP-gated nonspecific P2X7Rs on their cell membrane [9]. To investigate experimentally how P2X7Rs affect Ca^2+^ responses in these cells, a prolonged step pulse of 100 *µ*M ATP was applied at 60 s and Ca^2+^ signals from 99 ROIs were recorded. Ca^2+^ transients in these cells were also removed by simultaneously bathing cells in Hanks’ Balanced Salt solution. Despite the variability in the recorded Ca^2+^ responses, they consistently exhibited a rapid rise, characteristic of P2X7R activation (Fig. 5A). To better understand the differences between these response profiles, we clustered them into three distinct groups (Fig. 5B). The main feature that distinguished them from each other was the slow component following the fast rise. The slow component either increased the amplitude of the response with relatively steep gradient (top), increased it with a shallow gradient (middle) or gradually decreased it with a descending gradient (bottom) over time (Fig. 5B).

**Figure 5.**
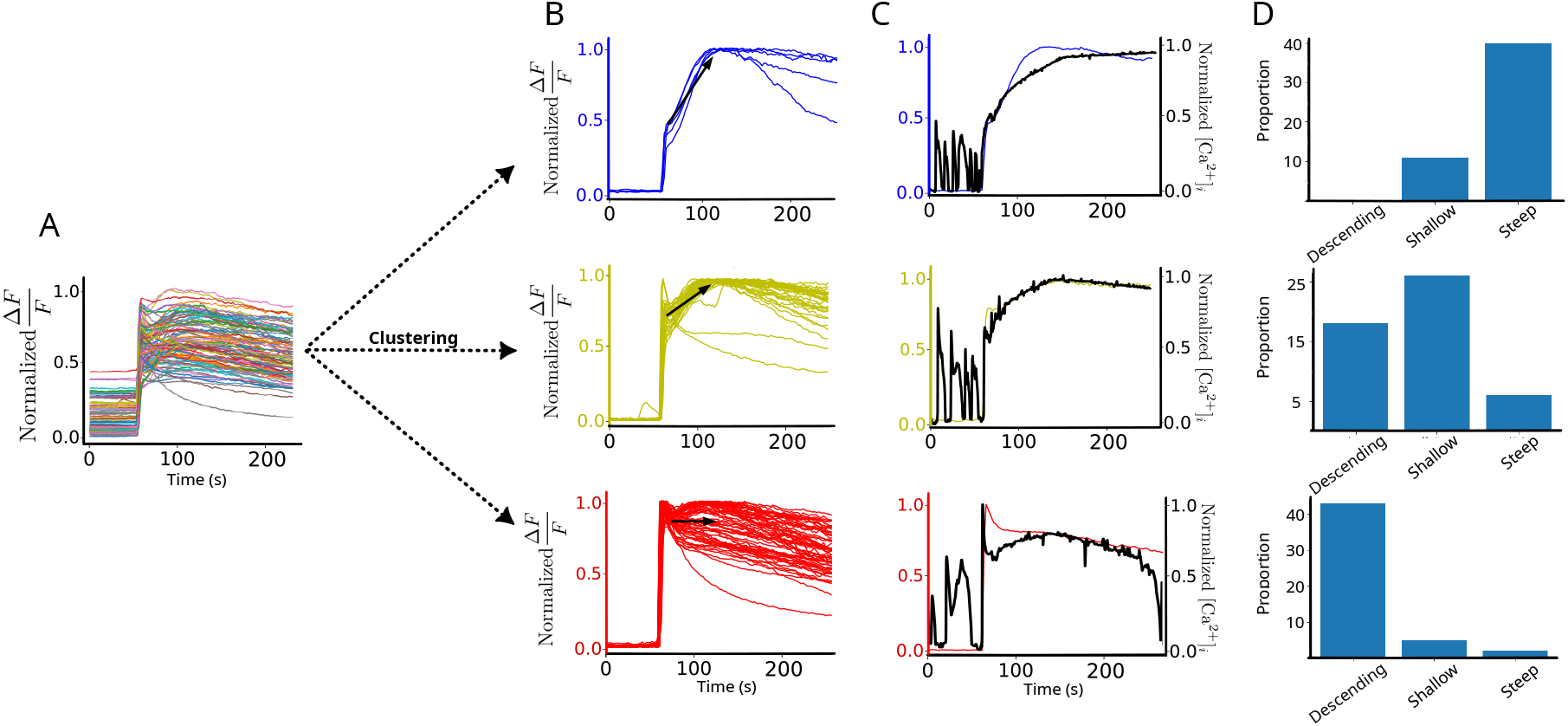
P2X7R-dependent Ca^2+^ responses in OPCs upon prolonged ATP stimulation. The set of all recorded 99 Ca^2+^ responses, normalized by their maximum, recoded in OPCs bathed in Hanks’ Balanced Salt solution and stimulated with 100 *µ*M ATP without interruption starting at 60 s. (B) The three distinct clusters of Ca^2+^ response profiles, including those with steep gradient = 0.009 (blue; top), shallow gradient = 0.04 (yellow; middle) and descending gradient = −0.001 (red; bottom). (C) Simulated Ca^2+^ responses (black) at different ATP concentrations: [ATP]= 0.1×10^−3^ M (top), 0.035×10^−3^ M (middle), and 0.028×10^−3^ M (bottom), overlaid on one representative Ca^2+^ recording from each cluster in B. Simulated responses were obtained by applying ATP for 180 s. (D) A set of 50 independent simulations of Ca^2+^ responses clustered into steep, shallow and descending gradient groups at different ATP concentrations: [ATP]= 0.1×10^−3^, (top) 0.05×10^−3^ (middle) and 0.028×10^−3^ (bottom). SCaLTs prior to ATP stimulation observed in the simulations in C are absent in the experimental recordings due to the bath solution.

To investigate how P2X7Rs contribute to the distinct response profiles of the three clusters, we performed simulations using the SSM with integrated P2X7R kinetics.. Because it is unclear how Hanks’ Balanced Salt solution removes SCaLTs (whether voltage-facilitated or intrinsic), we did not modify the model to block these random spiking events. Applying a prolonged step pulse of [ATP]= 0.1×10^−3^ M (i.e., 100 *µ*M) to the model generated responses that predominantly matched those obtained experimentally with a steep gradient and somewhat managed to generate a few responses with a shallow gradient, but failed to capture the descending gradient responses (Fig. 5C top, D top). Interestingly, by reducing the amplitude of the ATP step pulse to [ATP]= 50 *µ*M and 28 *µ*M, we were able to shift the Ca^2+^ responses produced by the SSM to predominantly match the experimentally observed profiles of the shallow (Fig. 5C middle, D middle) and descending gradient (Fig. 5C bottom, D bottom) responses, respectively. Taken together, these results suggest that P2X7Rs mediate the Ca^2+^ signals induced by prolonged ATP stimulation and that not all cells in culture experience the same [ATP] during these stimulation experiments. Moreover, the ability of the lower ATP concentrations to reproduce the spectrum of experimentally observed responses (Fig. 5C, middle and bottom; Fig. 5D, middle and bottom) suggests that stochasticity also plays a key role in shaping these responses.

### OPC Ca^2+^ Responses Induced by Glutamate

As is the case for P2X7Rs, OPCs also express the glutamatergic AMPARs and NMDARs [9]. To determine how glutamate affects Ca^2+^ signals in OPCs, we removed Ca^2+^ transients as before using the Hanks’ Balanced Salt solution and recorded from 100 ROIs in these cells when stimulated with prolonged step pulse of 100 *µ*M glutamate starting at 60 s (Fig. 6A). Despite exhibiting the same level of variability as that seen with ATP stimulation, the fast rise in Ca^2+^ responses upon glutamate stimulation was always a consistent feature across all responses. Clustering the recorded signals, as was done before, produced three distinct groups of responses distinguished from each other by the slow component following the fast rise. Once again, the slow component either increased the amplitude of the response with a steep gradient (top), increased it with a shallow gradient (middle) or gradually decreased it with a descending gradient (bottom) over time (Fig. 6B).

**Figure 6.**
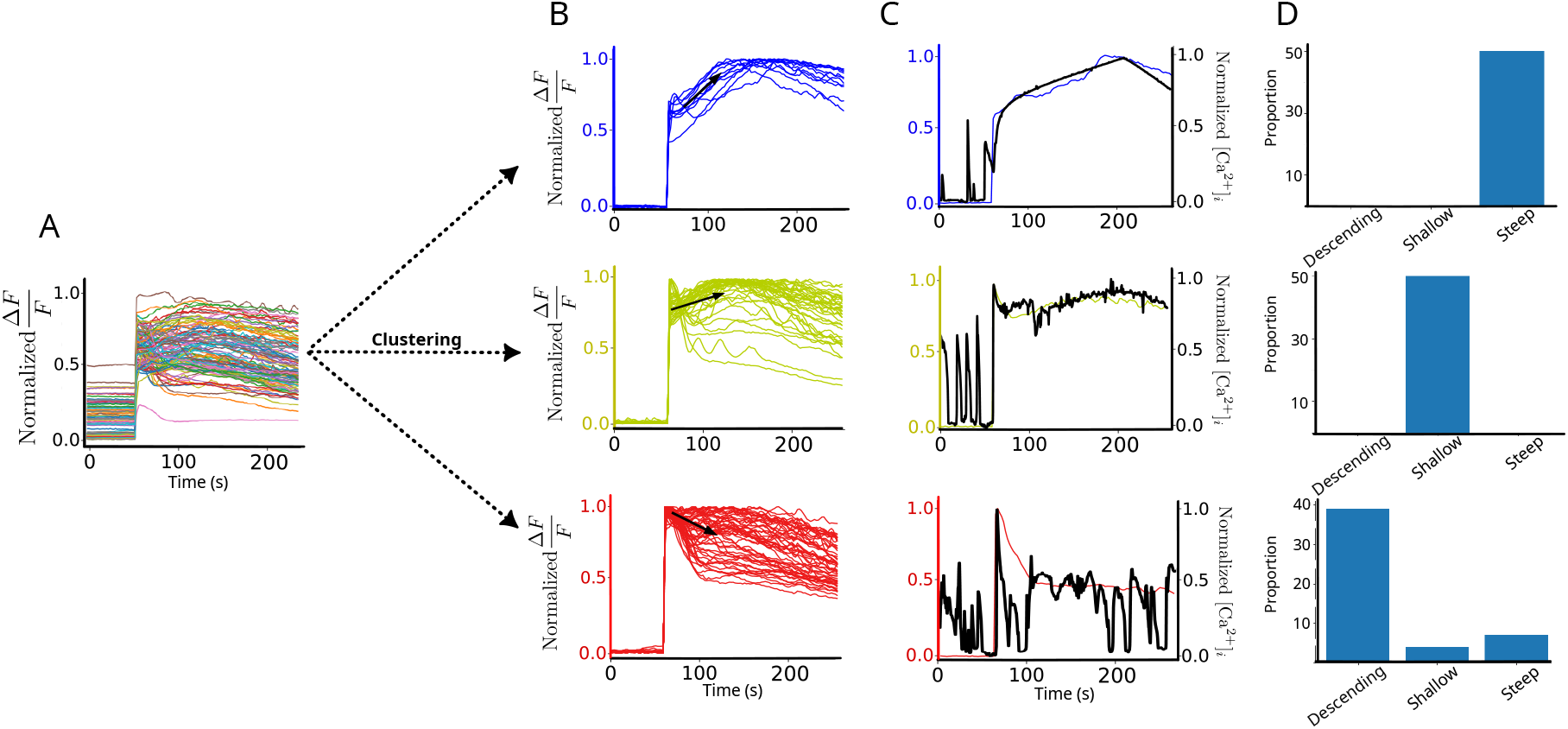
AMPAR and NMDAR-dependent Ca^2+^ responses in OPCs upon prolonged glutamate stimulation. (A) The set of all 99 recorded Ca^2+^ responses, normalized by their maximum, recoded in OPCs bathed in Hanks’ Balanced Salt solution and stimulated with 100 *µ*M glutamate without interruption starting at 60 s. (B) The three distinct clusters of Ca^2+^ response profiles, including those with steep gradient = 0.0081 (blue; top), shallow gradient = 0.0015 (yellow; middle) and descending gradient = −0.0047 (red; bottom). (C) Simulated Ca^2+^ responses (black) at different glutamate concentrations: [Glut]= 0.1 ×10^−3^ M (top) 0.04 ×10^−3^ M (middle), and 0.028 ×10^−3^ M (bottom), overlaid on one representative Ca^2+^ recording from each cluster in B. Simulated responses were obtained by applying glutamate for 180 s. (D) A set of 50 independent simulations of Ca^2+^ responses clusters into steep, shallow and descending gradient groups at different glutamate concentrations: [Glut]=0.1× 10^−3^, (top) 0.04 ×10^−3^ (middle) and 0.028 ×10^−3^ (bottom). SCaLTs prior to glutamate stimulation observed in the simulations in C are absent in the experimental recordings due to the bath solution.

Simulating the SSM, with the kinetics of AMPARs and NMDARs included, in response to 0.1×10^−3^ M (i.e., 100 *µ*M) glutamate stimulation, showed that it can closely but uniquely reproduce the steep gradient Ca^2+^ responses obtained experimentally (Fig. 6C top, D top). This was observed even in the presence of SCaLTs, which were not removed due to the unknown effects of Hanks’ Balanced Salt solution. To capture the shallow gradient responses (Fig. 6C middle; D, middle), we had to reduce [Glut] to 0.04×10^−3^ M. Further decreasing [Glut] to 0.028×10^−3^ M predominantly produced responses matching the descending gradient profile. Interestingly, some simulations at that lowest glutamate concentration also captured steep and shallow gradient responses. These results thus suggest that the glutamatergic receptors included in the model can reproduce the experimentally observed Ca^2+^ responses and that not all OPCs in culture experience the same glutamate levels. Finally, the findings once again highlight the crucial role of stochasticity in generating the wide spectrum of responses seen experimentally.

### Role of CICR in ATP- and Glutamate-Dependent Ca^2+^ Responses

One important and common feature of the Ca^2+^ responses induced by ATP and glutamate was the presence of an initial fast rise in Ca^2+^ signal upon stimulation. This feature appeared in all clusters of Ca^2+^ responses and in both conditions (Figs. 5A, B and 6A, B).

Interestingly, the entire Ca^2+^ response profiles (including the fast and slow components) associated with steep and shallow clusters associated with 100 *µ*M ATP stimulation were consistent with the P2X7R current profiles induced by high ATP (or its agonist BzATP) stimulation observed in previous studies [28]. Indeed, it was previously shown that the kinetics P2X7R model (Fig. 1F) alone is able to display two components in its current response upon high ATP stimualtion: an initial fast phase representing P2X7R channel opening followed by a slow rising phase representing P2X7R pore opening (or sensitization) [31, 30, 29, 28]. Incorporating this kinetic model into the SSM successfully reproduced the two phases of Ca^2+^ responses, suggesting that Ca^2+^ signaling is shaped by receptor-generated currents. However, when CICR was blocked in these simulations (i.e., by inhibiting fluxes through IP3R and RyR), the biphasic response disappeared entirely (Fig. 7A), leaving only a slow and gradual rise in [Ca^2+^]_i_ following ATP stimulation. This indicates that while ATP-dependent Ca^2+^ responses are influenced by P2X7R kinetics, the synergy between the fast dynamics of CICR and P2X7R-mediated influx is essential for generating the observed dynamics.

**Figure 7.**
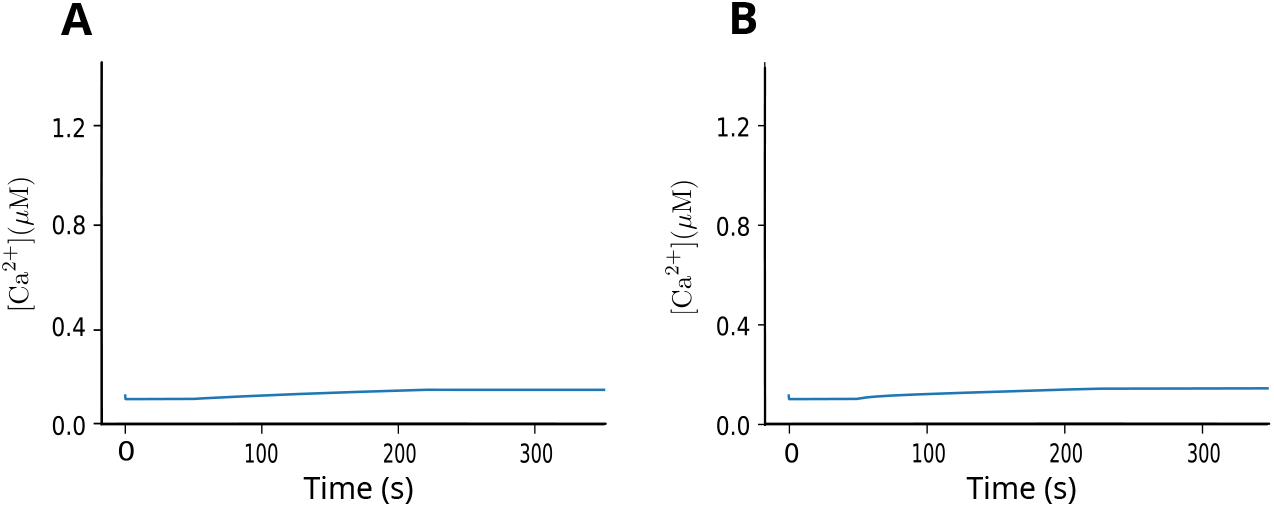
The stereotypical Ca^2+^ responses induced by ATP and glutamate in OPCs require Ca^2+^-induced Ca^2+^-release (CICR). Ca^2+^ responses triggered by 100 *µ*M (A) ATP, and glutamate stimulations for 180 s in the absence of IP3R and RyR fluxes. Notice the absence of pronounced responses seen in the simulated signals in Figs. 5C and 6C.

A similar effect was observed when simulating the SSM response to 100 *µ*M glutamate stimulation in the absence of CICR (Fig. 7A). In this case, the response also exhibited a slow, gradual rise, indicating that CICR is necessary for AMPARs and NMDARs to generate responses consistent with their kinetics. Collectively, these results underscore the critical role of IP3R- and RyR-mediated fluxes and their fast dynamics in enabling membrane currents to produce the experimentally observed Ca^2+^ responses.

### Ca^2+^ Responses to in vivo-Like Stimulation with ATP and Glutamate

The prolonged ATP and glutamate stimulations were performed *in vitro*, but such sustained stimulations do not occur *in vivo*. Instead, *in vivo* stimulations are more likely to be briefer and occur in a more stochastic manner, with varying amplitudes over time. It would therefore be interesting to investigate how Ca^2+^ signals respond to randomly timed ATP and glutamate stimulations, better reflecting physiologically-relevant *in vivo* conditions.

To address this, we applied random ATP and glutamate stimulations (Figs. 8A and 9A, respectively) in the form of brief 1 ms pulses of varying amplitudes to the SSM and quantified their effects on Ca^2+^ spiking (Figs. 8B and 9B, respectively). The ATP and glutamate pulses were delivered randomly at low (left), intermediate (middle), and high (right) frequencies (Figs. 8A and 9A, respectively). The resulting steady-state Ca^2+^ signals were then evaluated using two metrics: (i) the signal distribution, obtained by binning and averaging 50 simulated traces (Figs. 8C and 9C, respectively), and (ii) the full width at half-maximum of Ca^2+^ spikes encompassing both SCaLTs and evoked responses (Figs. 8D and 9D, respectively).

**Figure 8.**
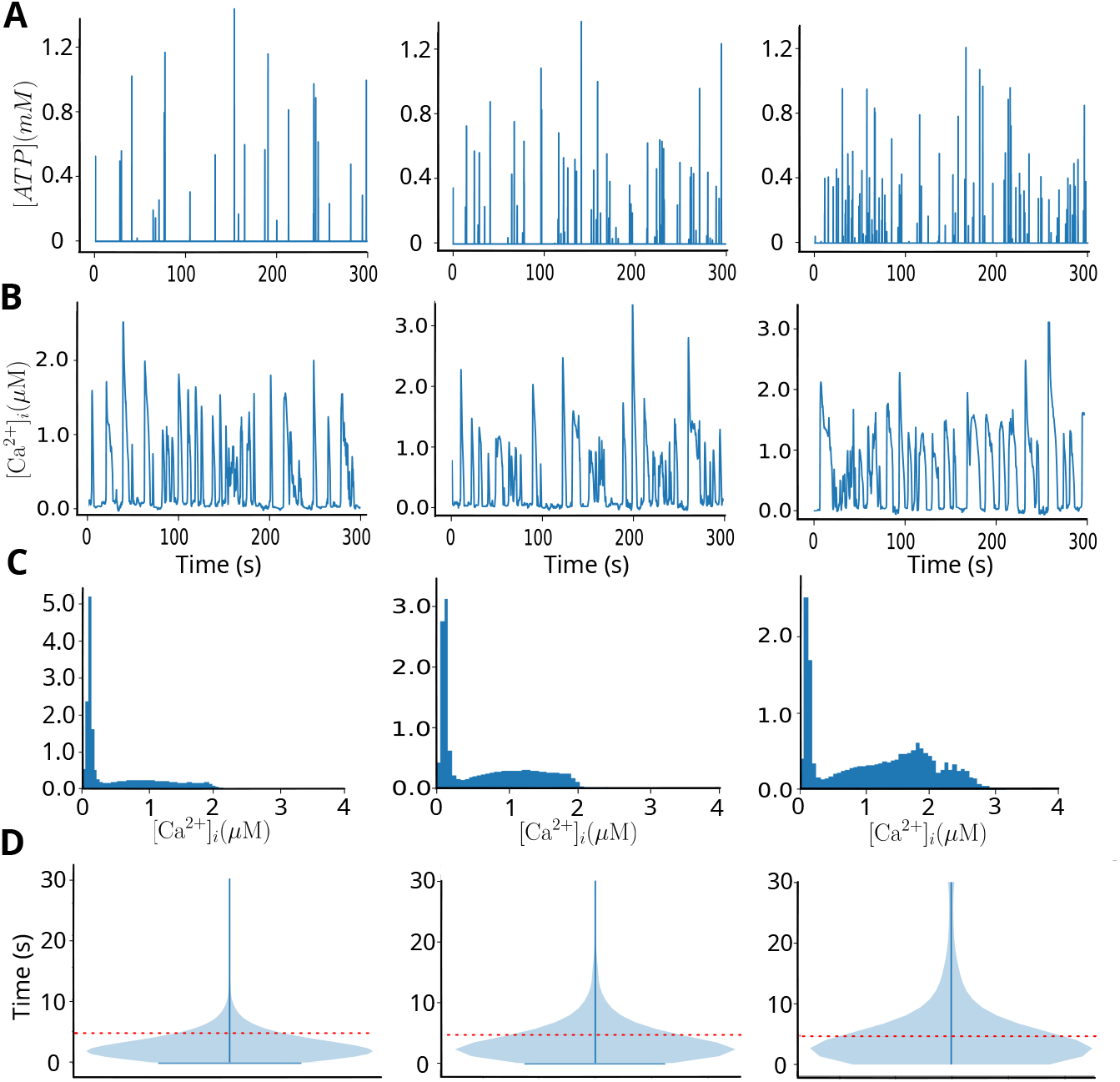
Simulated Ca^2+^ responses to random ATP stimulations mimicking *in vivo* conditions. (A) The random events of ATP stimulations in the form of brief 1 ms pulses with frequencies that follow a Binomial distribution whose mean is either low at 0.1 Hz (left), medium at 0.25 Hz (middle), or high at 0.5 Hz (right) and variance that is, respectively: 0.01, 0.1875 and 0.25. The amplitude of the ATP pulses are also random and follows a folded normal distribution with mean 0 and variance 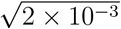. (B) Ca^2+^ transients (or spikes) combining both SCaLTs and evoked responses induced by the random ATP stimulations described in panel A, respectively. (C) Distributions of Ca^2+^ signals from 50 simulations of each of the three types of random stimulations shown in panel A, respectively, showing a significant increase in the amplitude and number of Ca^2+^ spikes. (D) Violin plots depicting the distribution of Ca^2+^ spike widths from 50 simulations of each of the three types of random stimulation shown in panel A, respectively. Red-dotted line separates wide spikes from narrow ones. Notice how an increase in the frequency of ATP stimulation leads to a moderate increase in the number of wide spikes.

Our results showed that random, brief ATP and glutamate stimulations induce a gradual shift toward elevated intracellular Ca^2+^ levels (Fig.8C) and an increase in the proportion of wider Ca^2+^ spikes (Figs. 8C and 9C, respectively) as stimulation frequency increases. As expected, more frequent ATP and glutamate applications led to prolonged elevations in Ca^2+^ signals, particularly when compared to control conditions. Although both ATP and glutamate induced similar effects, glutamate stimulation had a more pronounced impact, especially at low frequencies, suggesting that OPCs are more sensitive to glutamate than ATP (compare Fig. 8D, left to Fig. 9C, left). It is important to note that in both cases, the fluctuations in membrane voltage were not significantly affected by the frequency of these stimulations and remained roughly close to the resting membrane potential (Fig. S2).

**Figure 9.**
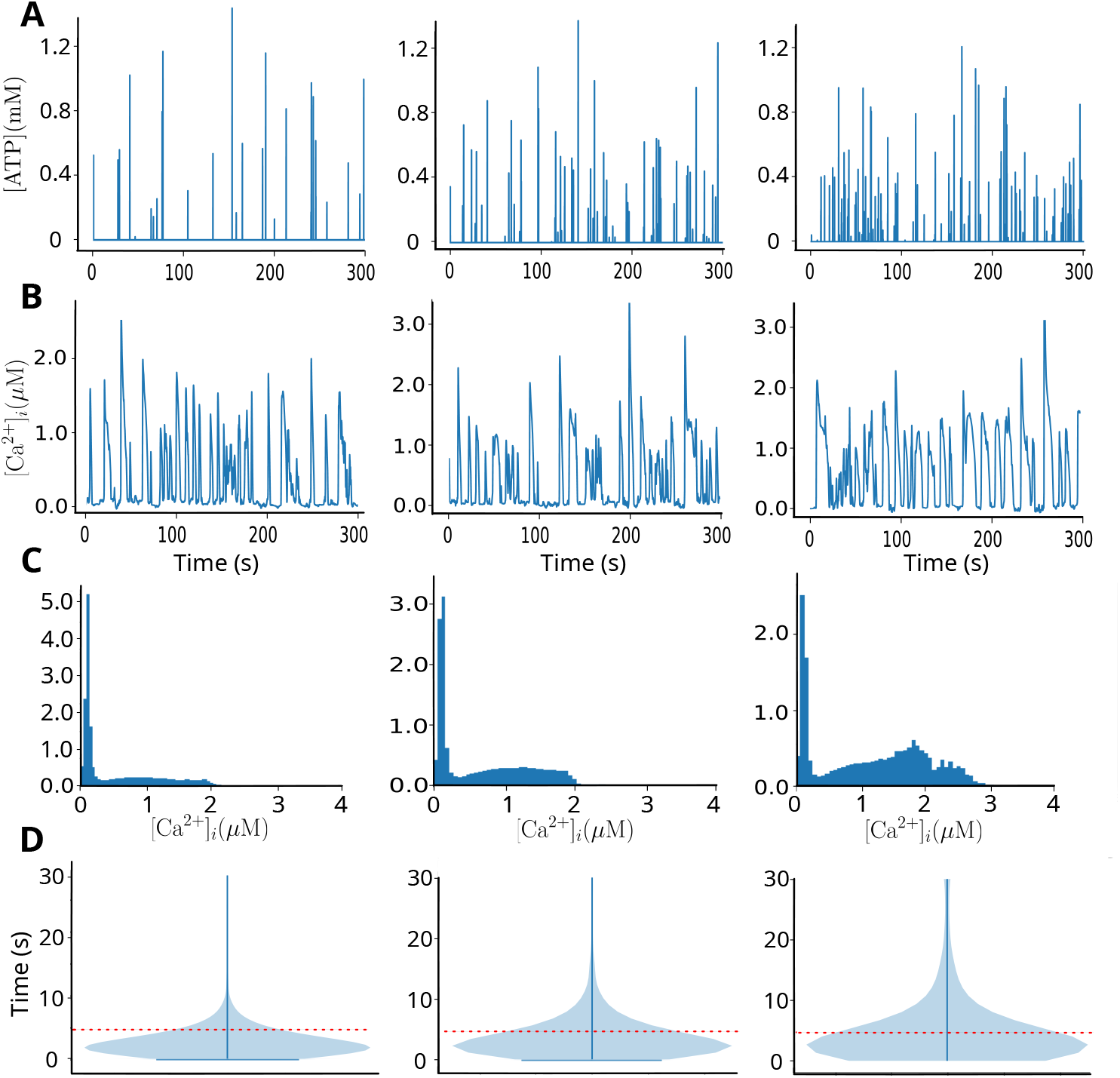
Simulated Ca^2+^ responses to random glutamate stimulations mimicking *in vivo* conditions. (A) The random events of glutamate stimulations in the form of brief 1 ms pulses with frequencies that follow a Binomial distribution whose mean is either low at 0.1 Hz (left), medium at 0.25 Hz (middle), or high at 0.5 Hz (right) and variance that is, respectively: 0.01, 0.1875 and 0.25. The amplitude of the glutamate pulses are also random and follow a folded normal distribution with mean 0 and variance 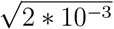. (B) Ca^2+^ transients (or spikes) combining both SCaLTs and evoked responses induced by the random glutamate stimulations described in panel A, respectively. (C) Distributions of Ca^2+^ signals from 50 simulations of each of the three types of random stimulations shown in panel A, respectively, showing a moderate increase in the number of spikes with large amplitudes without significant change in amplitude. (D) Violin plots depicting the distribution of Ca^2+^ spike widths from 50 simulations of each of the three types of random stimulation shown in panel A, respectively. Red-dotted line separates wide spikes from narrow ones. Notice how an increase in the frequency of glutamate stimulation leads to both a significant increase in the width of Ca^2+^ spikes as well as the number of such spikes.

Together, these findings indicate that ATP and especially glutamate stimulations mimicking *in vivo* conditions can induce broader Ca^2+^ spikes in OPCs; this may have significant implications on the functional role of OPCs and their maturation into oligodendrocytes.

## 4 DISCUSSION

Using a computational modeling approach, we investigated in this study how spontaneous and evoked Ca^2+^ responses due to stimulations with ATP and glutamate affect Ca^2+^ transients in OPCs. We expanded our previously developed spatiotemporal stochastic flux-balance Ca^2+^ model [12] to incorporate not only the dynamics of membrane voltage, modeled using the Hodgkin-Huxley formalism and regulated by L-type and T-type Ca^2+^ channels, but also the kinetics of both P2X7Rs and glutamatergic AMPARs and NMDARs. Because these latter receptors are not specific, we also tracked through the model the changes in Na^+^ and K^+^ concentrations ([Na^+^]_i_ and [K^+^]_i_, respectively). The resulting model both maintained the previously observed physiological and dynamic properties of the original model [12], as well as provided new insights into how VGCCs shape SCaLT dynamics and how purinergic and glutamatergic receptors drive evoked Ca^2+^ responses and modulate their spiking activity.

The two key components of the original model [12], including CICR via IP3Rs and RyRs and an Ornstein-Uhlenbeck noise process, enabled the model to previously generate random spiking events in the form of SCaLTs. In this study, the inclusion of VGCCs further ampli-fied these random spiking events and increased their frequency. Previous research [7, 8, 10] has suggested that myelination of axonal fibers is Ca^2+^-dependent and that the frequency of Ca^2+^ spikes plays a crucial role in regulating this process. Specifically, Ca^2+^ spike frequency must fall within an optimal “Goldilocks zone” to promote effective myelination by oligodendrocytes. The increase in spiking frequency induced by L- and T-type Ca^2+^ channels may help shift the system into this optimal range, thereby enhancing myelination. Further analysis of the steady-state dynamics of the DTM, in the absence of noise and diffusion, showed that stimulation with ATP and glutamate can transition the system into an oscillatory (spiking) regime. This transition favors deterministic dynamics over the stochastic nature of SCaLTs, potentially allowing Ca^2+^ spiking to stabilize within the so-called ‘Goldilocks zone’. P2X7Rs are well known purinergic receptors that, unlike P2×2Rs and P2×4Rs, exhibit two current phases upon stimulation with high [ATP] (or its agonist BzATP) [31, 30, 29, 28, 48, 49, 50]: a fast component (previously labeled *I*_1_), followed by a slow one (previously labeled *I*_2_). It was shown in these studies that the fast component is mainly due to channel opening, while the slow one is due to receptor sensitization. Recording the associated Ca^2+^ transients in these cells upon these ATP stimulations showed that evoked Ca^2+^ responses exhibited identical profiles with a fast rise (corresponding to *I*_1_) followed by a slow rise (corresponding to *I*_2_) in a manner similar to recordings shown in this study. By excluding CICR in the SSM, this two-phase profile disappeared completely, and only a very slow rise in [Ca^2+^]_i_ was observed. Interestingly, the tumor-derived GT1 cell line, commonly used to study gonadotropin-releasing hormone secretion, relies on CICR via IP3Rs to regulate intracellular Ca^2+^ concentration [51]. This mechanism enables GT1 cells to produce fast Ca^2+^ dynamics, allowing P2X7Rs to ultimately shape the Ca^2+^ responses observed experimentally [28] in a manner closely resembling the patterns reported in this study (Fig. 5).

Here, we highlighted the role of CICR in this process and emphasized that it extends also to glutamatergic receptors, including NMDARs and AMPARs.

Active axons communicate with local oligodendrocytes through two primary mechanisms: the vesicular release of glutamate and the non-vesicular release of intracellular ATP [52]. ATP is released via a volume-regulated anion channel, which is activated by osmotic swelling resulting from ion influx during action potentials [53]. Similarly, glutamate is released in an activity-dependent manner in both developing and mature oligodendrocytes [54]. Both ATP and glutamate regulate Ca^2+^ dynamics in myelinating oligodendrocytes, influencing myelin formation. Variations in the release patterns of these molecules along a single axon may contribute to the heterogeneous myelin distribution observed in the human cortex [55]. Approximately half of the Ca^2+^ transients in developing oligodendrocytes are driven by axonal action potentials, but the remaining ones occur independently of action potentials [10]. Interestingly, the positive correlation between the frequency of Ca^2+^ transients and speed of myelin sheath elongation is thought to be linked to the role of Ca^2+^ in actin polymerization in the myelin sheath [8]. Additionally, it has been experimentally demonstrated that these oscillations are activity-dependent, although the underlying mechanism remains unclear. In this study, we examined how applying random ATP and glutamate stimulation events mimicking *in vivo*-like conditions affect Ca^2+^ transients in the SSM. Our goal was to explore how Ca^2+^ spikes are formed as a combination of both evoked ATP- and glutamate-dependent responses and spontaneous events that are enhanced by VGCCs. Our results showed that even at low ATP and glutamate stimulation frequencies, the SSM generated broader Ca^2+^ spikes, which may serve a distinct role from narrower spikes, such as in cell differentiation. In contrast, the amplitude of the Ca^2+^ spikes remained largely unchanged. While this stability in amplitude aligns with previous studies suggesting that Ca^2+^ spike amplitude and shape are relatively insensitive to stimulation strength [56, 57, 58], the observed increase in spike width does not. This discrepancy is likely due to the stochastic nature of ATP and glutamate stimulation interacting with an excitable system, leading to random spiking events. More specifically, ATP and glutamate stimulation appear to drive the system from the spiking regime into an overstimulated regime [59], operating near the boundary between the two.

Stretch-activated Ca^2+^ channels (SACs) may contribute to Ca^2+^ influx in oligodendrocytes, particularly within the soma. These mechanosensitive channels are known to play a crucial role in cellular motility [60] and have been implicated in mechano-electric feedback mechanisms that regulate Ca^2+^ transients in cardiomyocytes [61]. In oligodendrocytes, osmotic swelling caused by Ca^2+^ influx could activate SACs, further amplifying Ca^2+^ currents. This feedback loop may play a role in regulating intracellular Ca^2+^ signaling and volume homeostasis in response to mechanical or osmotic changes. The role of SCA was not incorporated into the SSM but could be a promising avenue for exploring its impact on Ca^2+^ spikes.

Given that myelination in the central nervous system is closely linked to the neural activity of neighboring cells, our flux-balance-based SSM provided a framework for investigating the underlying dynamics of Ca^2+^ signals in OPCs. This approach offered fresh insights into the fundamental Ca^2+^ transients that drive neuro-oligodendrocyte signaling. Ultimately, the development of the SSM represents a crucial step toward unraveling the complexities of neuro-oligodendrocyte interactions, with potential implications for advancing our understanding of myelin-related processes.

## Supplementary Section

**Figure S1:**
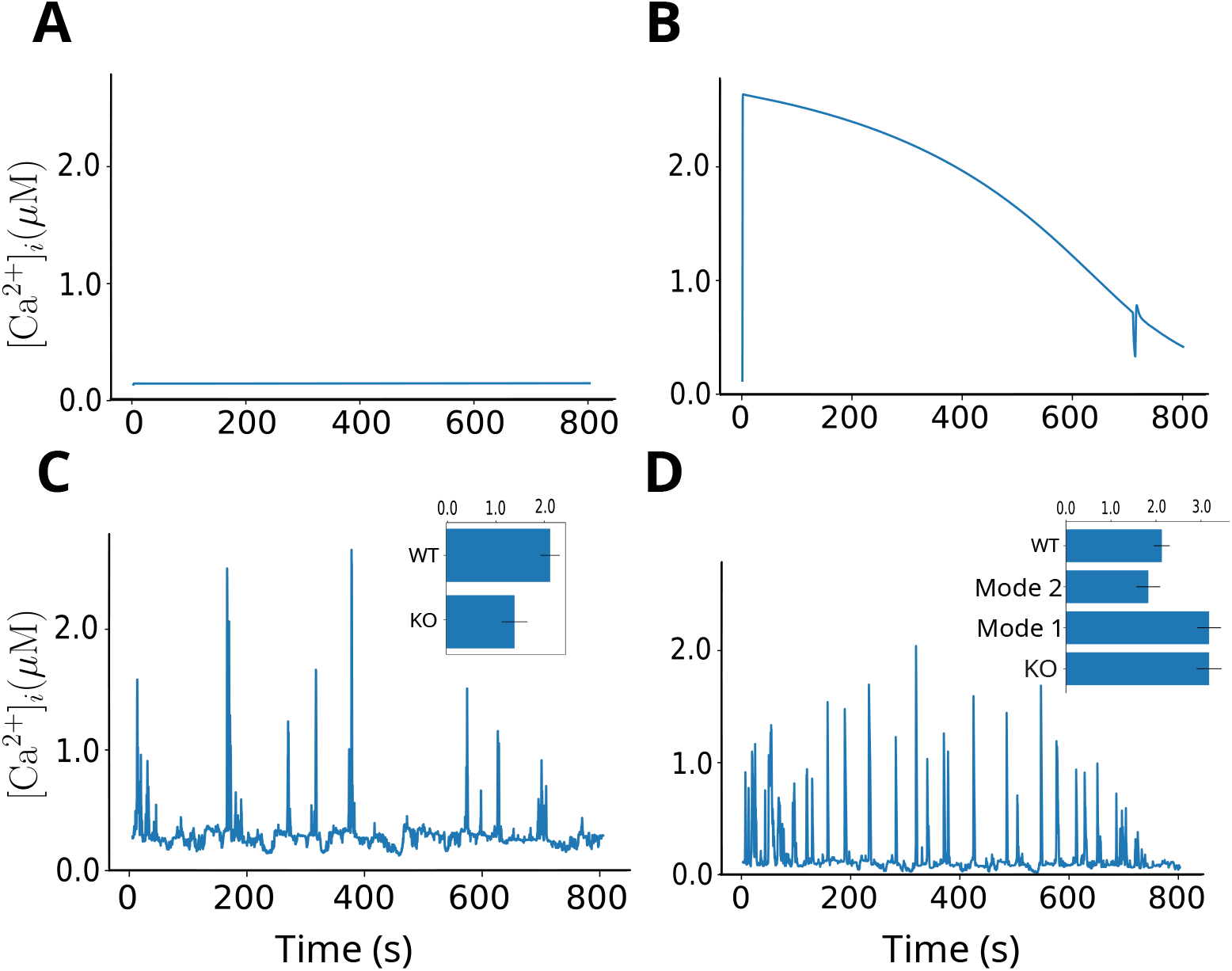
Effects of key Ca^2+^ fluxes on voltage-independent (or intrinsic) SCaLTs when the voltage, P2X7Rs, AMPARs and NMDARs are absent. As in [12], blocking (A) IP3Rs produces quiescent signal, (B) SERCA pumps produces one single Ca^2+^ transient, (C) RyRs produces reduced number of SCaLTs, and (D) mode 1, mode 2 and both modes of NCX produces a decrease, an increase and no effect, respectively. Insets in C and D are averages of 50 simulations for each condition.

**Figure S2:**
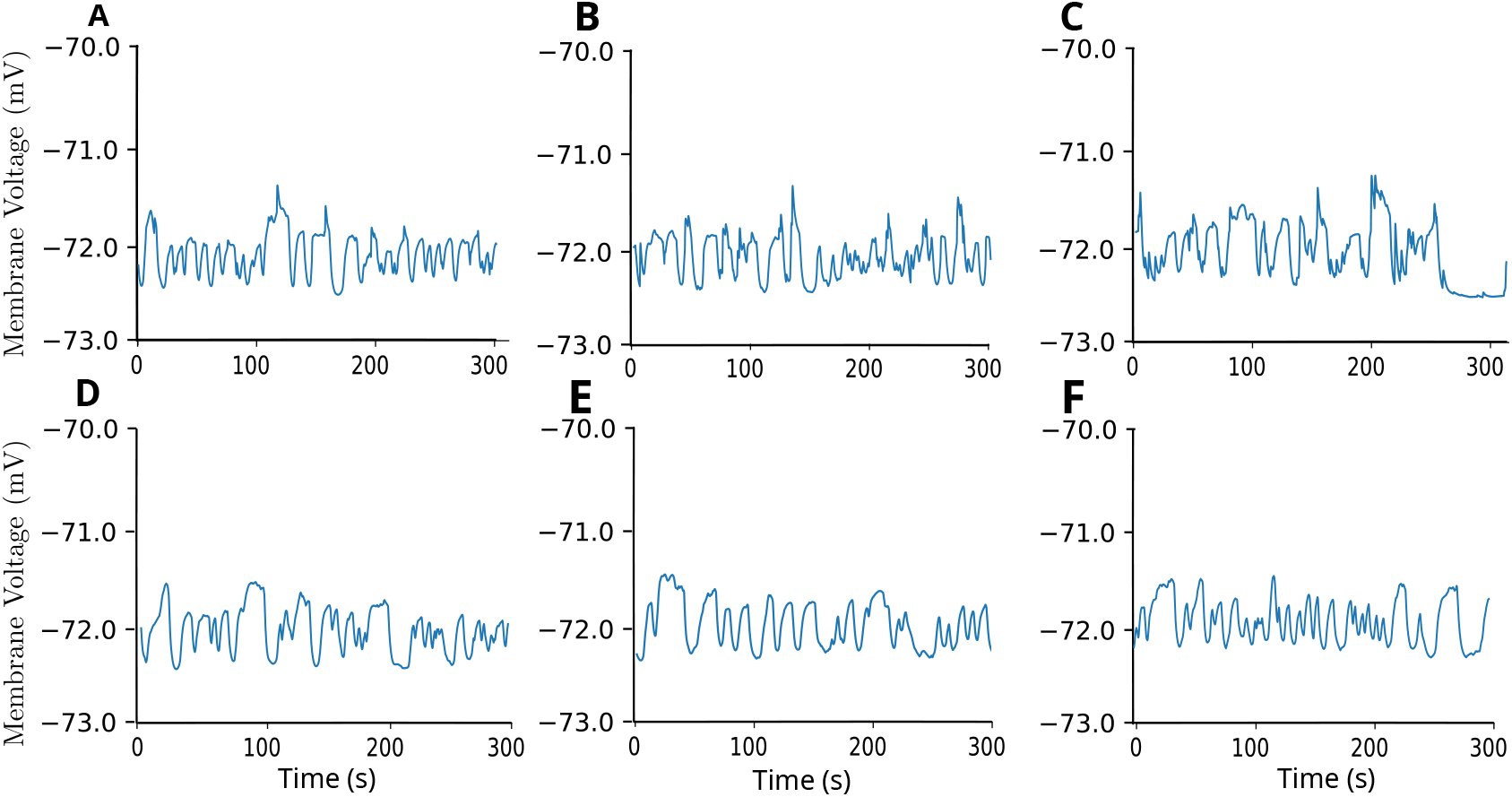
Fluctuations in membrane voltage upon random stimulations with ATP (top row) and glutamate (bottom row), mimicking in vivo conditions. Random stimulations are done at (A,D) low, (B, E) intermediate, and (C,F) high frequencies.

